# Examining Sleep Signals at the Cradle of Life: Can phylogenomic analysis of the Last Universal Common Ancestor (LUCA) reveal the fundamental role of sleep?

**DOI:** 10.1101/2025.03.21.644522

**Authors:** Seithikurippu R Pandi-Perumal, Konda Mani Saravanan, David Warren Spence, Sayan Paul, Ganesh Pandian Namasivayam, Saravana Babu Chidambaram

## Abstract

In common with most physiological activities, sleep is a highly evolutionarily conserved function. Nevertheless, the purpose of sleep remains inadequately investigated. One possible cause of this deficiency is the limitations of the traditional methods used for examining sleep. Up to this time, the mainstay tool used to look at the evolutionary basis of sleep has been phylogenetic analysis. This approach has provided many valuable insights into sleep, yet it has left many questions unanswered. The present study uses a relatively new hybrid technique at the interface of phylogenetics and genomics, known as phylogenomic analysis. This study is the first to use phylogenomic analysis to investigate the basis of sleep by evaluating the presence and conservation of sleep-related genes in the reconstructed genome of the Last Universal Common Ancestor (LUCA). Our gene set enrichment analysis of humans and LUCA indicates that the conserved sleep genes are linked to signaling, metabolism, and circadian rhythm pathways, suggesting that these genes possess primordial roles in essential physiological functions. These findings indicate that the component genes carry out essential physiological tasks that were subsequently repurposed to regulate sleep in more advanced organisms throughout evolution. This study lays the foundation for a systematic phylogenomic exploration of sleep-related genes, connecting molecular evolution with sleep science. By tracing the biological history of sleep to its deep evolutionary origins, our research presents novel insights into sleep’s nature, origin, and evolutionary function, paving the way for further interdisciplinary exploration of the biology of sleep.

**Graphical abstract:** 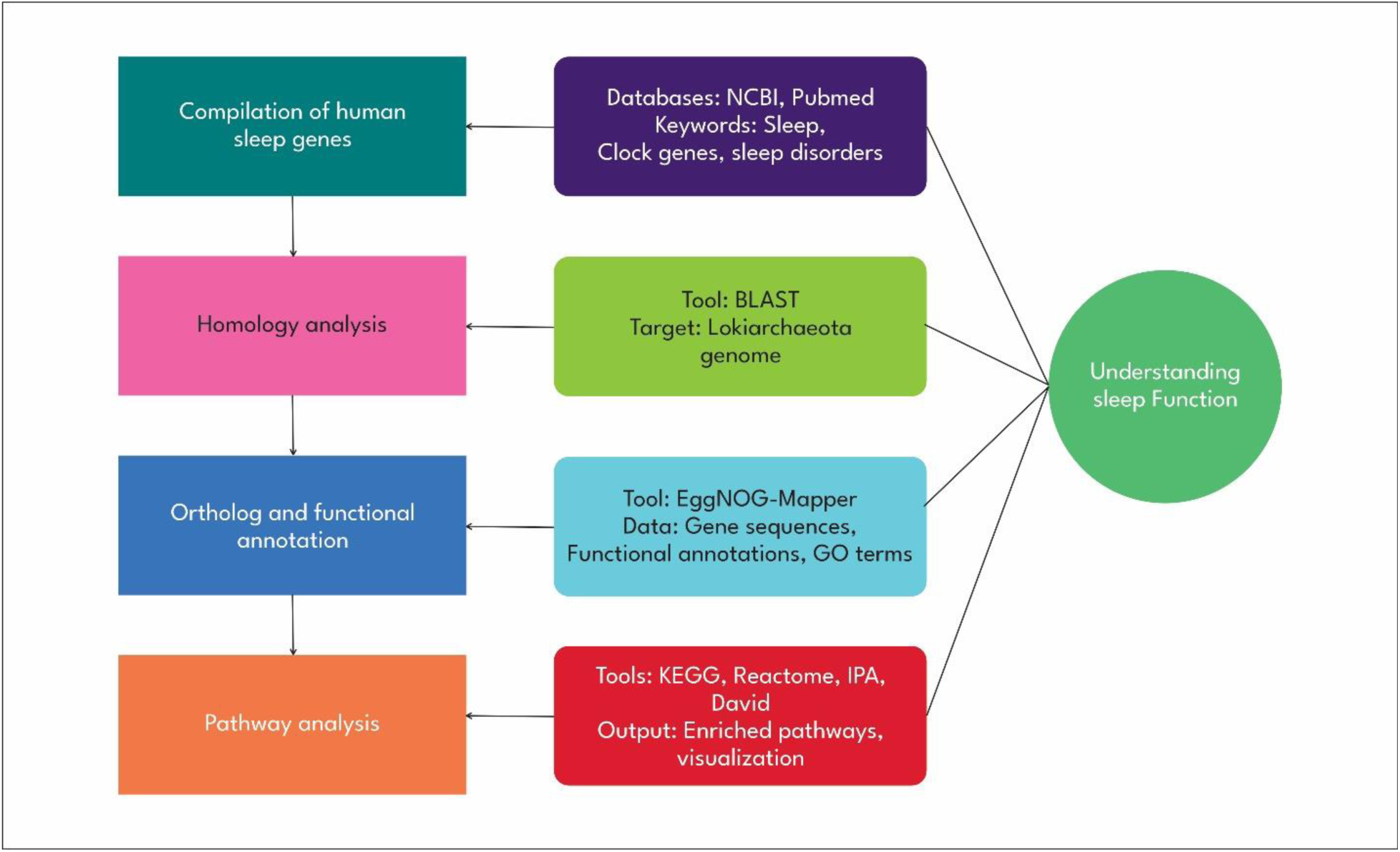

## Introduction

The genetics of sleep is a relatively nascent field. Over the last few decades, a few simple animal model organisms have helped to accelerate the field’s development [1]. These include but are not limited to, *Drosophila melanogaster* (fruit fly), *Caenorhabditis elegans* (nematode), and *Danio rerio* (zebrafish) [2].

Sleep is an active state documented across various organisms, ranging from simple invertebrates to complex mammals [3]. Several decades of research into sleep’s cause and function have suggested that the cnidarians, specifically the jellyfish Cassiopea, are most likely reported to be the oldest organisms to exhibit sleep-like behavior [4]. Despite their lack of a central nervous system, they display periods of quiescence or inactivity, behavioral traits that are consistent with sleep [5]. Cassiopea, for example, has sleep characteristics of higher animals, including reversible quiescence, Increased arousal threshold, and homeostatic control [4,6]. These findings suggest that sleep evolved much earlier than previously assumed, possibly more than 500 million years ago, in the common ancestors of cnidarians and bilaterians. This implies that sleep-like activity may be an ancient and fundamental biological mechanism deeply rooted in the evolutionary history of life on Earth [7].

Over the last few years, we have initiated a project to specifically investigate the phylogenomic aspects of sleep, which is viewed from the overlapping perspectives of phylogenetics and genomics [2,8,9]. This combined analytic approach has yielded results that have important implications for several sleep biology and medicine domains. By studying the genome sequences of various organisms to deduce their evolutionary links, phylogenomics provides insights that go well beyond typical phylogenetic investigations [8]. Essentially, phylogenomics advances our knowledge of the complexity and diversity of life while also improving our comprehension of evolutionary processes and offering several medical and biotechnological applications and environmental conservation.

First utilized by Darwin [10], the “Tree of Life” (TOL) is one of the key explanatory models used today in the field of evolutionary biology. One of the most striking processes that the model highlights is how incredibly sophisticated the most well-known modern living forms are. They vary in morphology, physiology, behavior, ecology, and other life strategies. In general, these organisms look and behave differently from one another. This conclusion, one in which the differences between modern organisms are more salient than their similarities fails to take an evolutionary perspective into account, i.e., it tends to see the twigs and branches of the tree rather than its foundational roots. It is argued here that more is to be learned from how organisms are the same rather than from how they are different. The view taken in the present study is that it is only possible to understand the core processes of complex functions by investigating the earliest beginnings of these biological processes, i.e., from the standpoint of evolutionary biology.

A reasonable starting point in analyzing the roots of sleep would be a study of our shared biological ancestors. To some extent, this effort recapitulates the history of the study of evolutionary biology itself. The last universal common ancestor (LUCA or the progenote) is believed to have originated some 4.2 billion years ago [11]. Its adaptive processes reflected the stressful conditions that often prevailed at the time. The LUCA thrived in harsh, gassy, metal-laden, extremely hot plumes from seawater reacting with lava that erupted through the ocean floor, known as deep-sea hydrothermal vents.

The LUCA, a hypothetical (theoretical construct) anaerobe and a hyper-thermophile to mesophilic, is reconstructed based on comparing the genomes of currently living organisms [12]. The three domains of life presently on Earth, bacteria, archaea, and eukaryotes, all share universal genes (UG) and are thought to have descended from LUCA, the most recent common single-cell ancestor [13]. It is further believed that LUCA’s earliest bifurcation point on the TOL was the fundamental divergence recognized as Archaea and Bacteria. The implication of this process, which is that all organisms are related, was a foundational template for evolutionary biology as it developed in the 19^th^ century. Naturalists Jean-Baptiste Lamarck (1744-1829) and Charles Darwin (1809–1882) initially proposed that all species shared a common origin, an idea that became integrated with the study of genetics. The postulate of the last universal common ancestor (LUCA) emerged because of scientists’ efforts to decipher life’s genetic sequences and biochemical processes in the following century [14]. This work provided insight into the history of evolution.

According to scientists, the LUCA had a small genome with astounding complexity. Its genome consisted of about 2.5 million bases (2.49–2.99 Mb) and encoded around 2,600 proteins, a biological richness that gave it the adaptability required for survival in its harsh geochemical habitat [11].

Many researchers in sleep science believe that sleep serves an ultimate purpose, a tenet based more on logical inference than hard evidence. This assumption, however, has not enjoyed universal acceptance, and many have pointed to the longstanding need for basic research on this issue. Phylogenomic analysis could be a means to investigate this matter further. This is what we have tried to do in this study, wherein we mapped the genes linked to sleep and sleep disorders with a synthetic genome such as LUCA. We mapped the genes linked to sleep and sleep disorders with phylogenetically remote species such as LUCA and its early variants.

## METHODS

### COLLECTION OF SLEEP GENES

A systematic compilation of genes related to sleep, its functions, and sleep disorders was made from the NCBI gene search. This produced a lengthy list of sleep-related terms such as clock, circadian, sleep, circadian rhythm, insomnia, sleep disorder, sleep regulation, and sleep-related genes. These terms were further analyzed for their broader significance in sleep research. This second-level analysis applied filters such as ‘Homo sapiens’ to focus on human-related genes, along with additional filters such as ‘publication date ranges’ and ‘research type’ to align the study with pertinent research objectives. Gene summaries analyzed alongside relevant literature for each outcome, focusing on annotations, facilitated the identification of direct one-to-one involvement with sleep processes or disorders. The participation of these elements in sleep function was confirmed by cross-referencing the list with academic papers utilizing NCBI’s PubMed database [15]. The assembled dataset included 280 sleep-associated genes. Each gene was linked to a PubMed citation that outlined its role in sleep, as demonstrated by experimental studies in animal models. Supplementary resources, including GeneCards and Online Mendelian Inheritance in Man (OMIM), were utilized to comprehensively validate the genes and assemble new reviews on sleep research [16]. A meticulously curated collection of genes acquired using this analytical approach facilitated the elucidation of the functions of sleep genes. The objective was to provide a foundation for our subsequent phylogenomics research and analysis.

### COLLECTION OF LUCA GENES

This approach outlines the method for identifying genes associated with LUCA utilizing the NCBI Gene database [17]. The search query included the terms “core conserved genes” and/or “universal genes,” indicating the user’s intent to identify evolutionarily conserved and potentially ubiquitous genes throughout all domains of life. Filters for genomic, protein-coding, and annotated genes narrowed the results to solely functional and conserved genetic elements. The search results yielded a total of 861 genes. The results table includes details for each gene, including its Gene ID, description, chromosomal location, aliases, and MIM number, which can substantiate the findings. This search approach demonstrates the systematic application of specific criteria to the database, including protein-coding and annotated genes, together with keyword filtering. This lists genes potentially associated with LUCA or other conserved functions. The vast database enables researchers to refine their search parameters to identify candidate genes for more in-depth evolutionary or functional studies.

### BLAST ANALYSIS OF SLEEP GENES AGAINST LUCA GENOME

The complete gene sequences of LUCA were obtained using the procedure described above. The standard protocol was applied to do a BLAST (Basic Local Alignment Search Tool) analysis of sleep genes against the LUCA genome [18,19]. The nucleotide or protein sequences of the discovered sleep-related genes were manually assembled from the National Center for Biotechnology Information (NCBI) in FASTA format. The NCBI BLAST homepage (URL: https://blast.ncbi.nlm.nih.gov/Blast.cgi) was accessed. The desired database was specified by opting for the LUCA genome stored as a local database. The human sleep gene sequences were fed into the BLAST interface by uploading them. Subsequently, the search parameters were adjusted, including the E-value threshold, match/mismatch scores, gap penalties, and word size, to perform the BLAST search. This systematic approach permitted critical insights into the evolutionary preservation and functional resemblances of genes associated with sleep across human and LUCA genomes.

### GENE ONTOLOGY ENRICHMENT AND PATHWAY ENRICHMENT ANALYSIS

The enrichGO function was utilized from the clusterProfiler package in R to identify the biological processes, molecular activities, and cellular components associated with the gene set of interest [20]. The gene list, comprising functionally relevant genes in the LUCA genome, was created utilizing their Entrez identification numbers. Gene Ontology term annotations were acquired from the most recent version of org.Hs.eg.db (or an analogous package for the relevant organism). The investigation was conducted independently for the three GO domains. This annotation system is categorized into three distinct classifications: Biological Process (BP), Molecular Function (MF), and Cellular Component (CC). The over-representation test determined the enrichment of a particular GO word, and the likelihood of a gene-term relationship was evaluated using the hypergeometric distribution model. The False Discovery Rate (FDR) threshold of <0.05 was utilized to identify the most enriched GO keywords. The results were subsequently illustrated using dot plots and bar plots generated by the clusterProfiler program [20].

An enrichment analysis of the pathway was also conducted to determine the role of the gene set in biological pathways. This was done using the Kyoto Encyclopedia of Genes and Genomes (KEGG) pathway database and the enrichKEGG function of clusterProfiler [21]. Gene Entrez numbers were used, and KEGG data for species such as humans was used for data selectivity. Significant enriched pathways were determined by performing a hypergeometric test, and any pathway with an adjusted probability (FDR) of less than 0.05 was considered significant. To achieve high accuracy of the results, the gene universe was defined as all the genes expressed in the dataset or genome of interest. The results of the pathway diagrams and the enrichment scores were drawn in dot plots, bar plots, and network plots.

### INTEGRATION AND VISUALIZATION SOFTWARE

An integrated study compared enrichment analysis results for GO keywords and KEGG pathways. The clusterProfiler program generated enriched word networks and enrichment maps to identify biological processes and pathways that share common and related activities [20]. For visualization, enriched terms and routes were illustrated utilizing tools such as ggplot2, employ, and cent plot [20] within the cluster profile. These annotations provided insight into the gene set’s biological importance and functional co-regulation.

All analyses were performed using R programming language utilizing the clusterProfiler and org.Hs.eg.db (or species-specific database) packages [20]. The FDR < 0.05 was employed as a criterion for significance, along with additional settings such as qvalueCutoff and pvalueCutoff, which are recommended by default in the clusterProfiler software [20].

## RESULTS

### HUMAN SLEEP GENES IN THE LUCA GENOME

This comparative study aimed to analyze the evolutionary conservation of human sleep genes in the genome of the Last Universal Common Ancestor (LUCA). It explored the conservation of 59 sleep-regulating genes, encompassing transcription factors, enzymes, receptors, and structural proteins across several species. The study elucidated the biological importance of these genes, indicating that their physiological functions may be essential for numerous biological processes and that forces of evolution may influence gene expression [22]. **Table 1** presents a comprehensive list of 59 genes, descriptions, lengths, e-values, and mean similarities (sim mean). These data illustrate the diversity of proteins examined, including transcription factors, enzymes, receptors, and structural proteins, while acknowledging the multitude of tasks proteins serve in biological processes. All entries’ stated e-value is zero, indicating that these alignments are highly significant. This means the 59 sleep-related human genes were conserved with the reference LUCA genome used for comparison. Glutamine synthetase, MHC class II antigens, and Sirtuin 1 exhibit remarkable conservation across species, indicating their potential critical physiological functions, as evidenced by their sequence similarities of 99.80%, 99.85%, 99.57%, and 99.77%, respectively. The low similarity in proteins such as Titin (48.74%) and Hypoxia-inducible factor 1-alpha (53.22%) may be attributed to adaptive or interspecies variations.

**Table 1.**
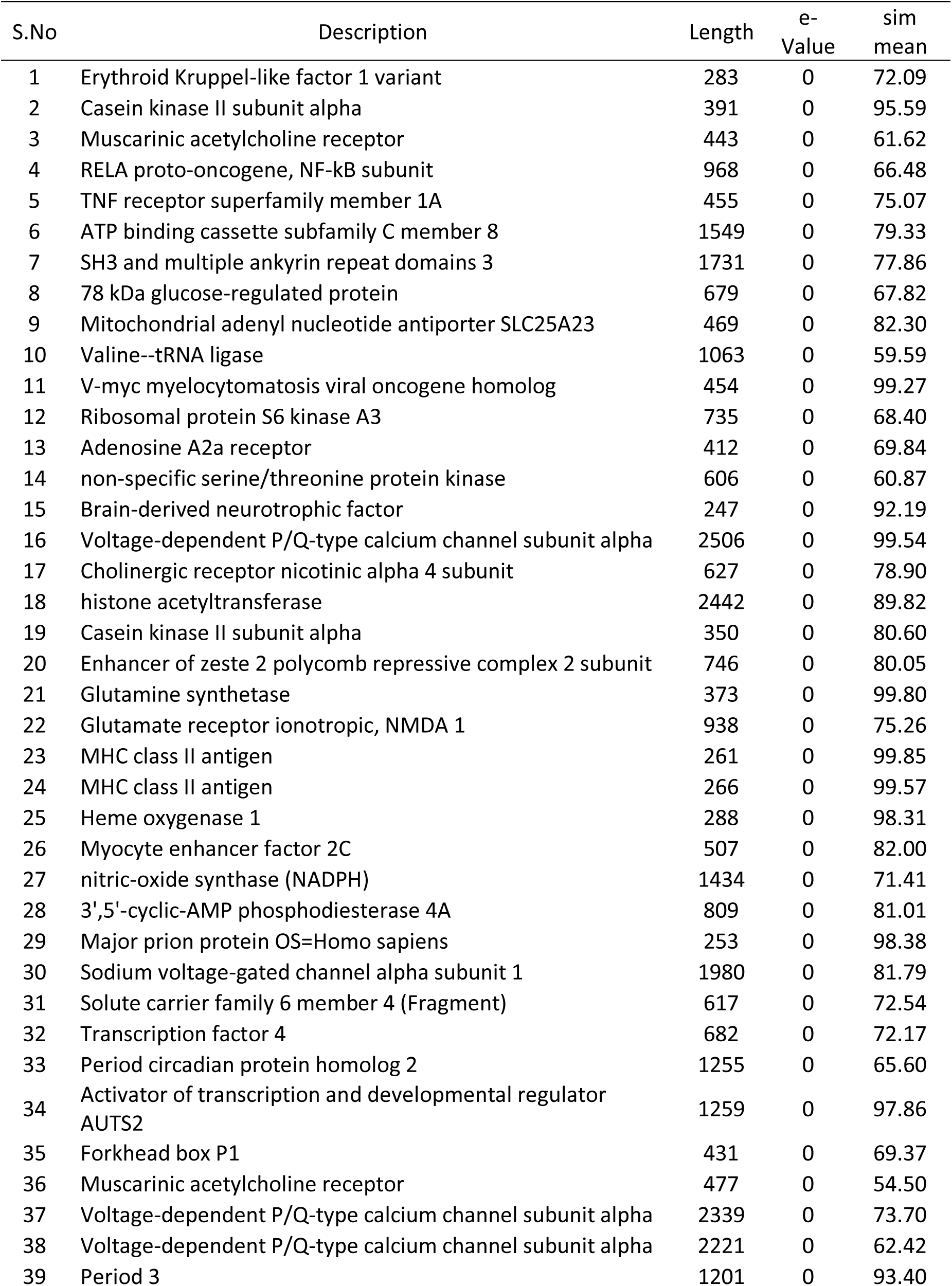
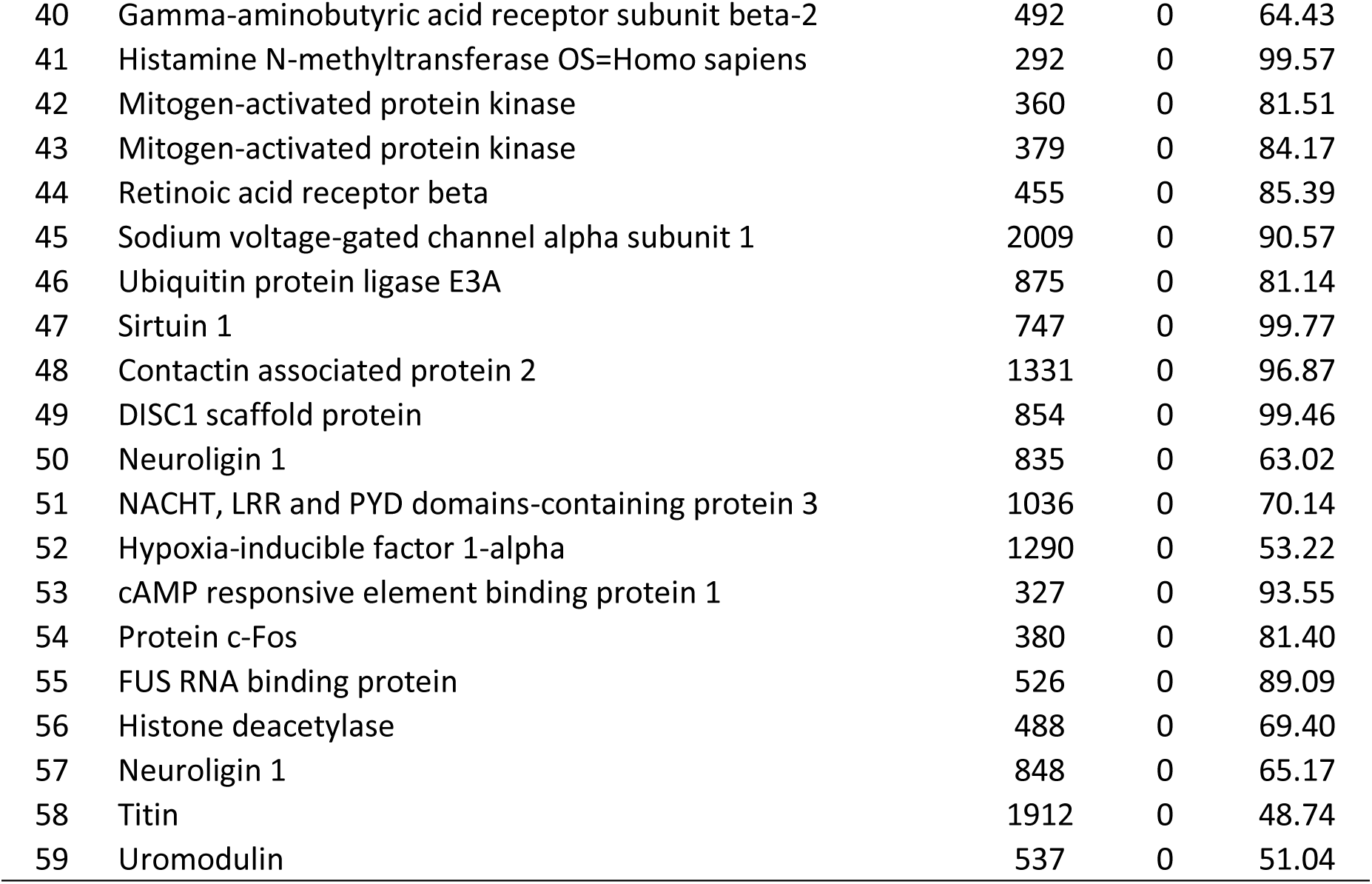
A comprehensive list of sleep gene orthologs present in LUCA, including their description, length, e-value, and mean similarity (sim mean).

### GENE ONTOLOGY ENRICHMENT ANALYSIS BIOLOGICAL PROCESSES

The Gene Ontology (GO) enrichment analysis illustrates biological processes pertinent to the examined genes. The processes are organized by GeneRatio (the ratio of input genes to each phrase) and the corrected p-value. The most enriched GO processes consist of genes associated with regulating nervous system development, circadian rhythm, and muscle system operations, which are essential physiological functions of the body (**Figure 1**). The circadian rhythm and its regulation serve as input genes that underpin the connection between sleep-wake cycles, with the circadian regulation of sleep being a conserved mechanism. Furthermore, heart contraction, blood circulation regulation, and nervous system development enhancement indicate the link between sleep-related genes and cardiovascular and neurological systems. Supplementary components that regulate synapse polarity and dendritic development suggest their involvement in brain plasticity and cognitive activities governed by sleep.

**Figure 1.**
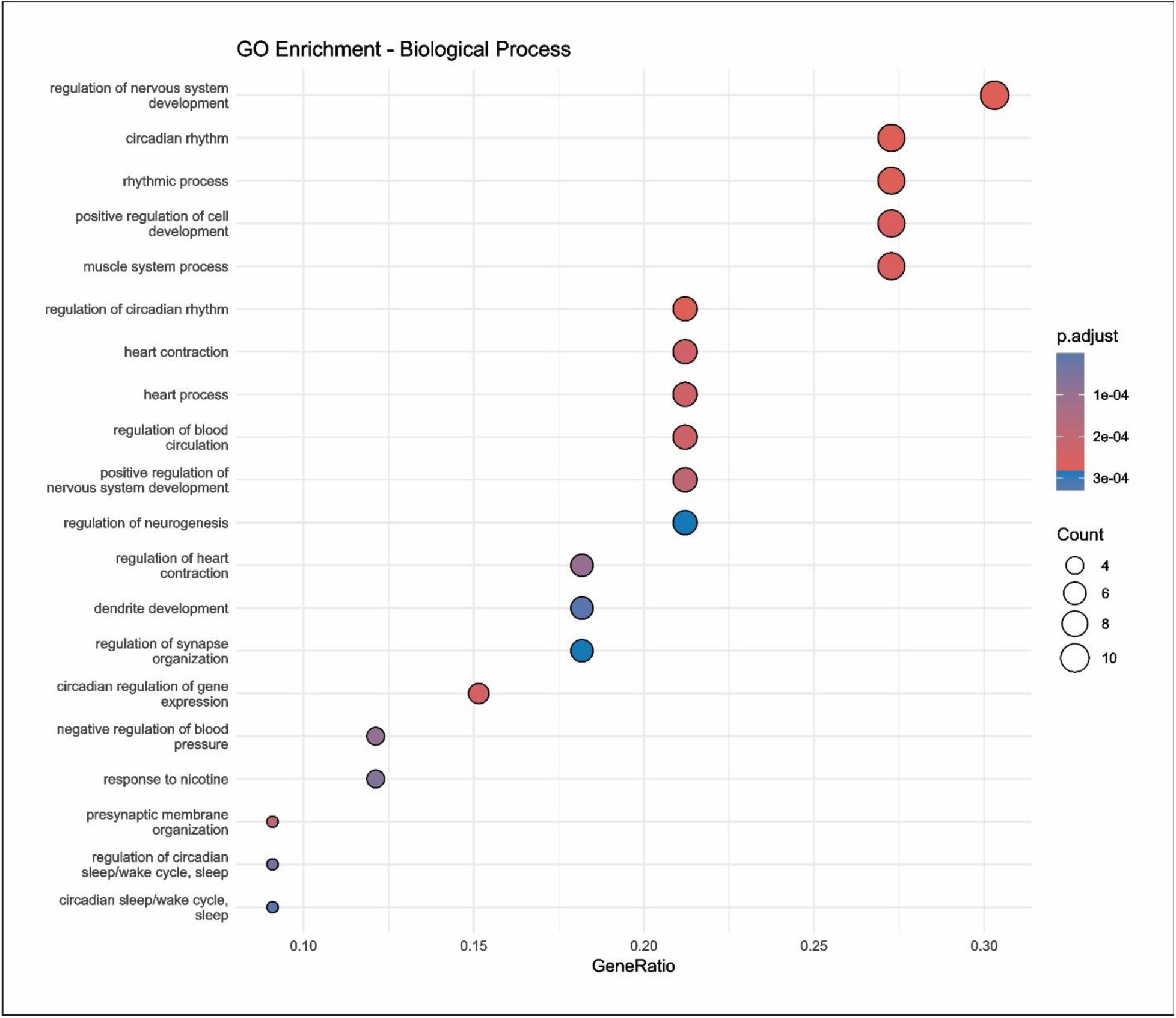
Gene Ontology (GO) enrichment analysis for biological processes associated with sleep-related genes. The x-axis represents the GeneRatio, while the y-axis lists enriched biological processes. The dot size indicates the number of genes involved, and the color gradient reflects statistical significance (adjusted p-value), with red representing the most significant terms.

Gene-environment interactions or physiological reactions to external stimuli may be deduced from responses such as nicotine response and negative blood pressure control. Finally, the processes related to the circadian sleep/wake cycle connect these genes to sleep biology. The diameter of the dots signifies the number of genes linked to the term; the dimensions of the circles are proportional to gene involvement in the enriched terms, while the color gradient (blue to red) reflects the p-value, with red denoting the most significant term. This depiction reinforces the association of these genes with brain, cardiovascular, and circadian systems, linking them more closely to sleep and its associated biological processes.

### MOLECULAR FUNCTION

The Gene Ontology (GO) analysis for biological activities offers crucial insights into the roles of sleep-related genes. The horizontal axis denotes GeneRatio, indicating the percentage of genes associated with specific molecular functions depicted on the vertical axis **(Figure 2)**. The dot size reflects the gene quantity, while the color gradient indicates the adjusted p-value, with darker hues representing elevated p-values. Other notable molecular functions include DNA-binding transcription factor activity, transcription coregulator activity, and RNA polymerase II transcription coactivator activity. The description of these roles emphasizes the capacity of sleep genes to modulate the transcription of other genes that participate in various cellular activities. Additionally, the Gene Ontology (GO) Molecular Function keywords chromatin DNA binding and histone deacetylase binding suggest an epigenetic dimension to the findings, implying that sleep-associated genes may influence chromatin architecture and transcriptional repression.

**Figure 2.**
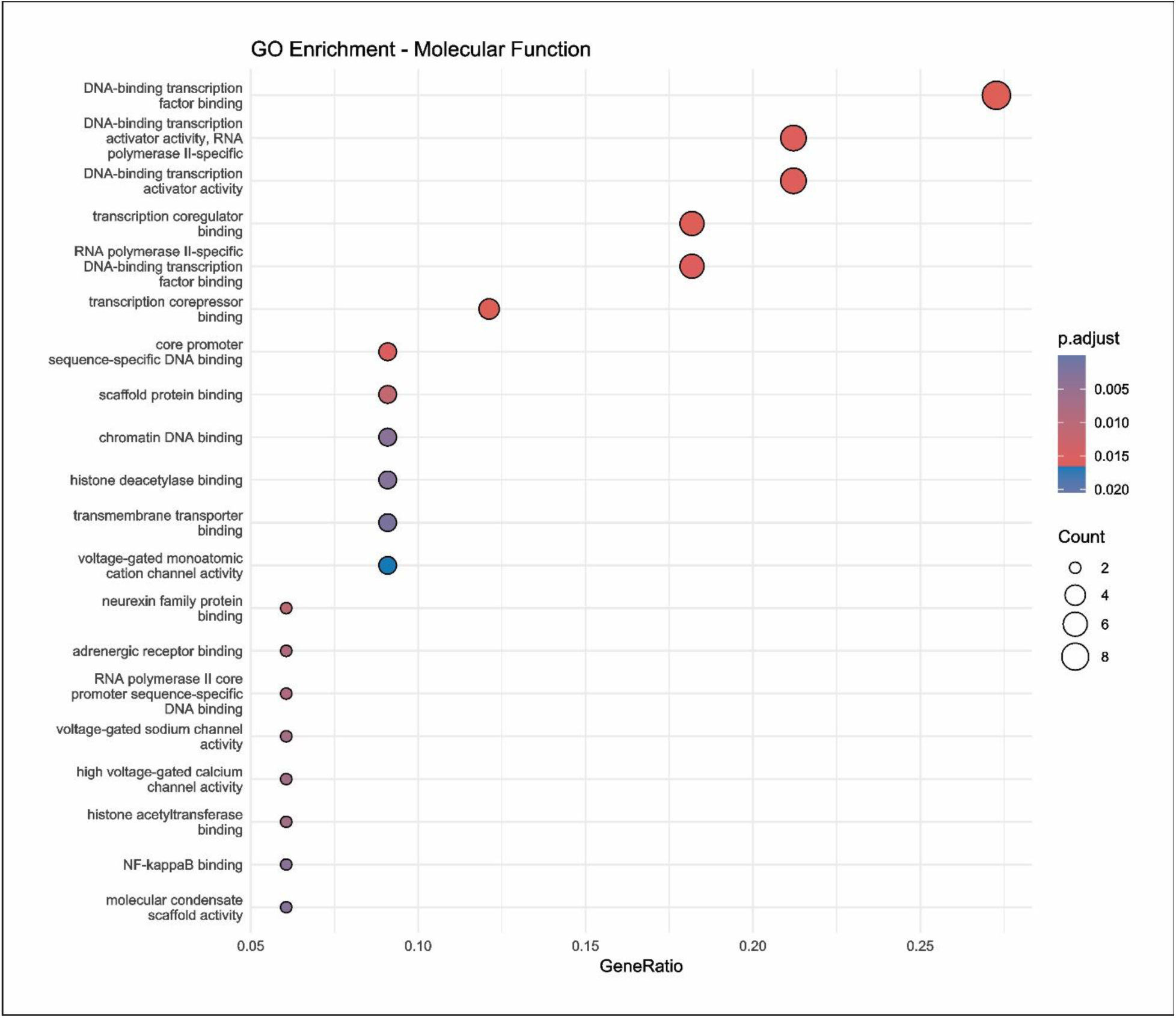
GO enrichment analysis of molecular functions associated with sleep-related genes. The x-axis represents the GeneRatio, the y-axis lists enriched molecular functions, and dot sizes indicate the number of genes involved. The color gradient reflects statistical significance (adjusted p-value), with darker shades indicating higher significance.

The activity of monoatomic cations, sodium channels, and voltage-gated ion channels further substantiates the involvement of sleep genes in neuronal signaling and the modulation of membrane potential, which are essential for circadian rhythm and sleep-wake cycle regulation. Scaffold protein binding and molecular condensate scaffold activity suggest their role in protein-protein interactions and the assembly of macromolecular complexes potentially implicated in sleep regulation. Adrenergic receptor binding and nuclear factor kappa-B (NF-κB) binding are significantly linked to stress responses and immunological modulation, suggesting that sleep genes integrate environmental and organismal signals. The studies reveal that sleep genes are implicated in several molecular activities, including transcriptional control, ion transport, and protein scaffolding, which are essential for sleep regulation.

### CELLULAR PROCESSES

The GO analysis of the cellular component highlights sleep-related genes’ localization and potential functions inside specific cellular compartments **(Figure 3)**. The most enriched components are neuronal synapses, such as neuron-to-neuron synapses, glutamatergic synapses, GABAergic synapses, and associated regions of the synaptic membrane, presynaptic membrane, and postsynaptic membrane. These enhancements underscore the importance of sleep genes in synaptic transmission and plasticity, which are essential for memory consolidation and neuronal stability during sleep [23]. Additional enhancements encompass the cation channel complex and voltage-gated calcium and sodium channels, pivotal in ion flux and action potential generation. Their engagement with sleep mechanisms reflects the importance of ion mobility in neuronal excitability and the rhythmicity of circadian clocks. Moreover, complexes such as the ESC/E(Z) and PcG proteins, which are involved in chromatin remodeling and gene expression, illustrate the connection between epigenetic control and sleep.

**Figure 3.**
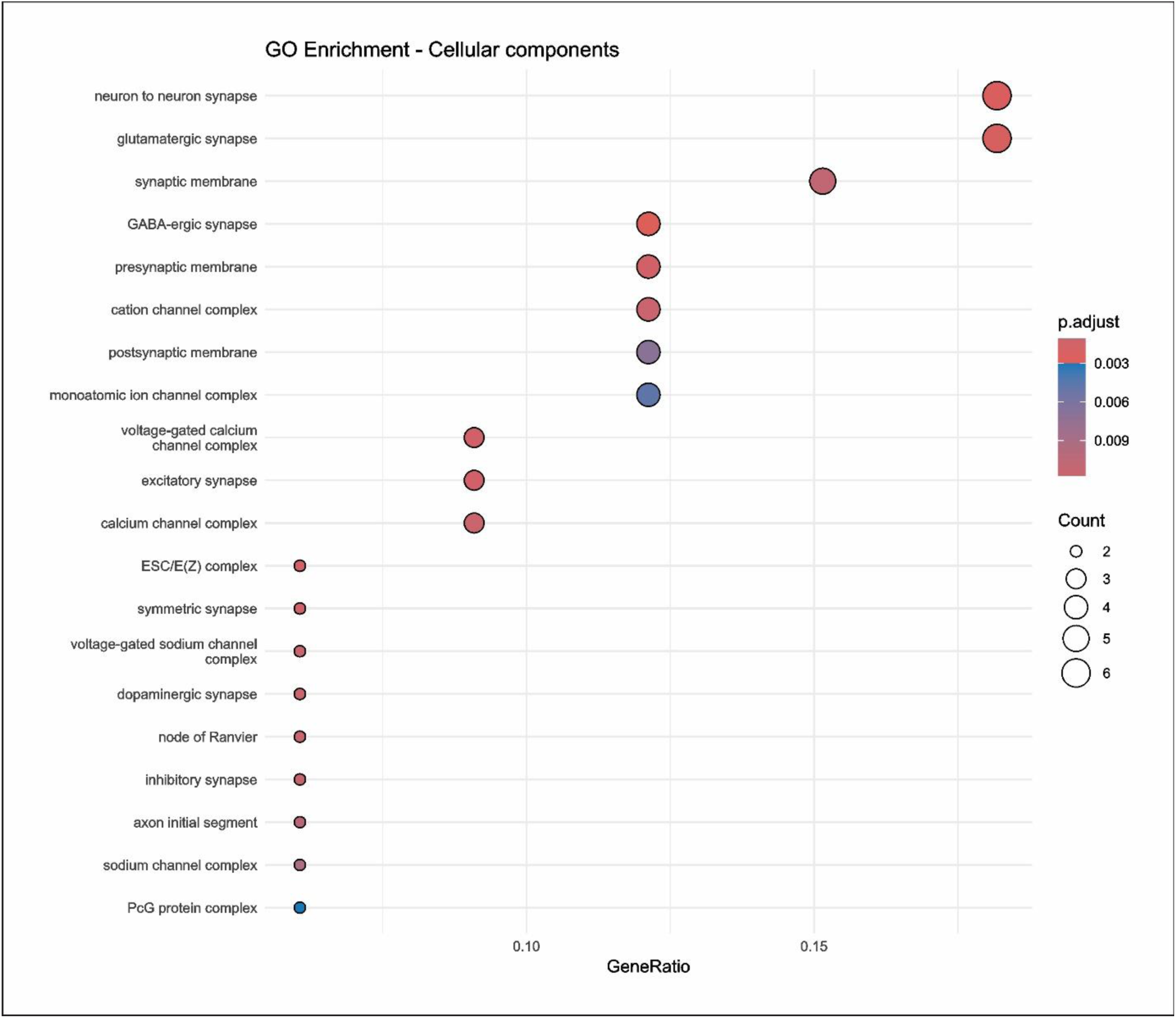
GO enrichment analysis for cellular components associated with sleep-related genes. The bubble plot shows enriched cellular components, including synaptic structures (e.g., neuron-to-neuron synapse, synaptic membrane) and ion channel complexes (e.g., voltage-gated calcium and sodium channel complexes). Bubble size represents gene count, while color indicates the adjusted p-value, with red denoting the most significant terms. These components emphasize the role of sleep-related genes in neuronal signaling, synaptic plasticity, and ion flux regulation.

An overview of dopaminergic and inhibitory synapses highlights the role of neurotransmitter systems in sleep regulation, particularly with the equilibrium between excitatory and inhibitory systems. The improvement of structures such as the AIS and the node of Ranvier, which are essential for signal transmission, underscores the significance of neuronal architecture and connectivity in maintaining sleep/wake cycles. The findings highlight the intricate interplay between sleep-associated genes and neural signaling, synaptic plasticity, and epigenetic processes.

### PATHWAY ENRICHMENT ANALYSIS

#### KEGG PATHWAY ENRICHMENT

Figure 4 displays the list of sleep-related genes alongside the KEGG pathway enrichment analysis pertinent to these genes. The most enriched pathways are the cAMP signaling route, circadian rhythm entrainment, and the circadian rhythm pathway, underscoring the significance of these genes in regulating the sleep-wake cycle and cellular signaling. Sleep-associated genes have been observed to influence pathways essential for cellular signaling in response to stimuli and neurotransmitters, including the MAPK signaling pathway and dopaminergic synapse. The increase of pathways associated with chemical carcinogenesis and microRNAs in cancer indicates a potential link between sleep regulation and cancer processes via hormone or stress mechanisms. The advent of novel addiction-related pathways, specifically cocaine and amphetamine addiction, further substantiates the role of these genes in the control of brain plasticity and reward signaling. The data underscores the multifaceted role of sleep-associated genes in regulating circadian rhythms, intercellular signaling, and other physiological and pathological processes. These pathways have potential therapeutic uses in sleep disorders and other associated diseases when addressed.

**Figure 4.**
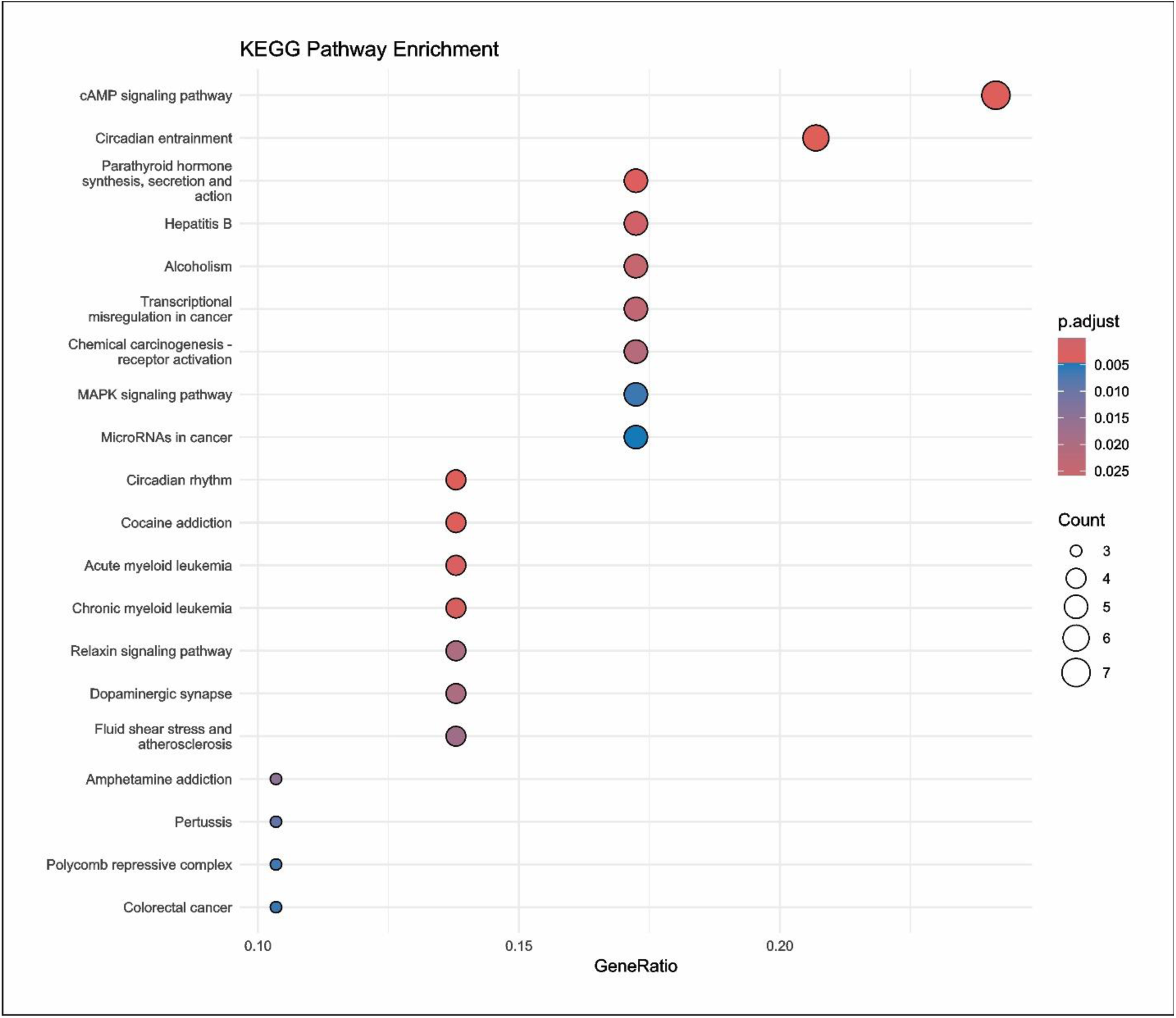
KEGG pathway enrichment analysis of sleep-related genes. Significantly enriched pathways include cAMP signaling, circadian entrainment, and MAPK signaling, highlighting the roles of these genes in circadian regulation, cellular signaling, and neurological functions. Pathway significance is indicated by adjusted p-values (color gradient), gene ratio (x-axis), and gene count (circle size).

### REACTOME PATHWAY ENRICHMENT

Figure 5 illustrates a Reactome Pathway Enrichment study, revealing pathway groups considerably overrepresented in a gene or molecular dataset. The routes are depicted along the y-axis, while the x-axis illustrates the GeneRatio, indicating the proportion of genes associated with the pathways of interest concerning the total number of input genes. The area of the circles correlates with the number of genes linked to a specific pathway, while the color indicates the p.adjust, reflecting the significance level of enrichment. Upon ascending the plot, an examination of the GeneRatio reveals those pathways such as “MAP kinase activation” and “Interleukin-17 signaling” exhibit elevated values, indicating their robustness within the data. These pathways regulate immunological responses, inflammation, and cellular communication. The enhancement suggests that the gene set may be involved in immune control and inflammatory signaling pathways. Several TLR signaling pathways are significantly elevated, specifically TLR 10, TLR 5, TLR 7/8, TLR 9, and TLR 3. These pathways are essential to the innate immune system as they enable cells in the body to identify infections and respond suitably. The identification of both MyD88-dependent and MyD88-independent pathways indicates the presence of several signaling systems within the dataset.

**Figure 5.**
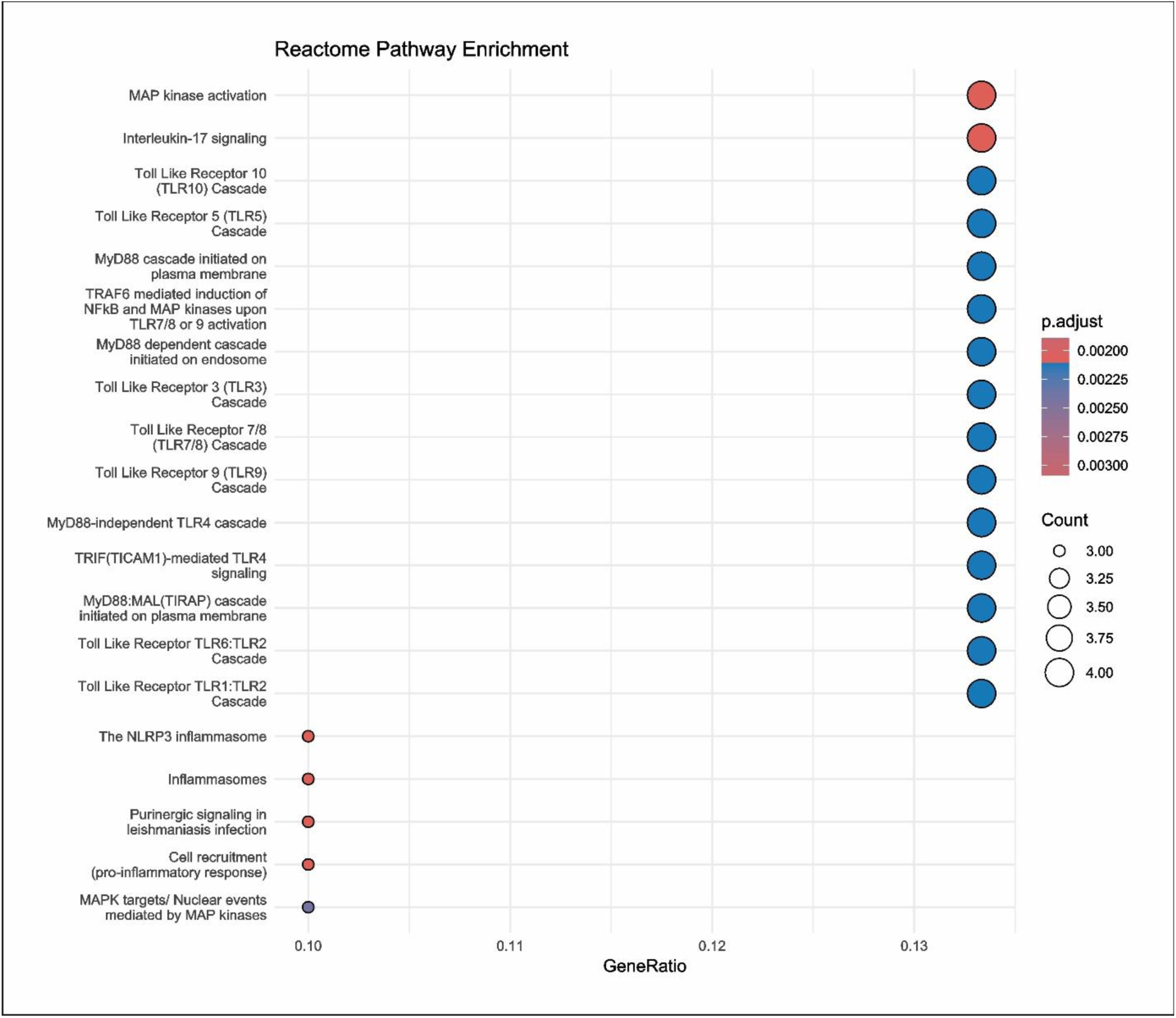
Reactome Pathway Enrichment analysis showing significantly enriched pathways. The x-axis represents the GeneRatio, while the y-axis lists the pathways. Circle size corresponds to the gene count, and color indicates the adjusted p-value (p.adjust), with darker shades reflecting higher statistical significance. Key pathways involve immune signaling, inflammation, and pathogen recognition.

Additional pathways such as ‘The NLRP3 inflammasome’ and ‘Inflammasomes’ are also enriched, indicating a potential role of inflammasome signaling in the observed biological activity. The current findings may indicate processes related to inflammation or pyroptosis. Furthermore, “Purinergic signaling in leishmaniasis infection” and “Cell recruitment (pro-inflammatory response)” denote specific pathogen-associated immune responses and cellular motility. The p.adjust values indicate the magnitude of these findings, with smaller p-values (e.g., red and dark orange circles) denoting greater statistical significance in enrichment.

### PROTEIN FAMILIES’ ENRICHMENT

Figure 6 illustrates the twenty protein families and domains most significantly overrepresented in the sample. The y-axis represents the protein families and domains, while the x-axis indicates the quantity of proteins within each specific family or domain. The highest-ranked families and domains, such as “ voltage-gated Na^+^ channel (VGSC), Cytoplasmic Domain,” “Voltage-Gated Cation Channel Calcium (CaV) and Sodium (NaV),” and “Voltage-Dependent Channel Domain Superfamily,” unequivocally demonstrate a significant overrepresentation of ion channel proteins. The results suggest focusing on membrane transport and electrical signaling routes, crucial for neural communication, muscular contraction, and signal transduction pathways.

**Figure 6.**
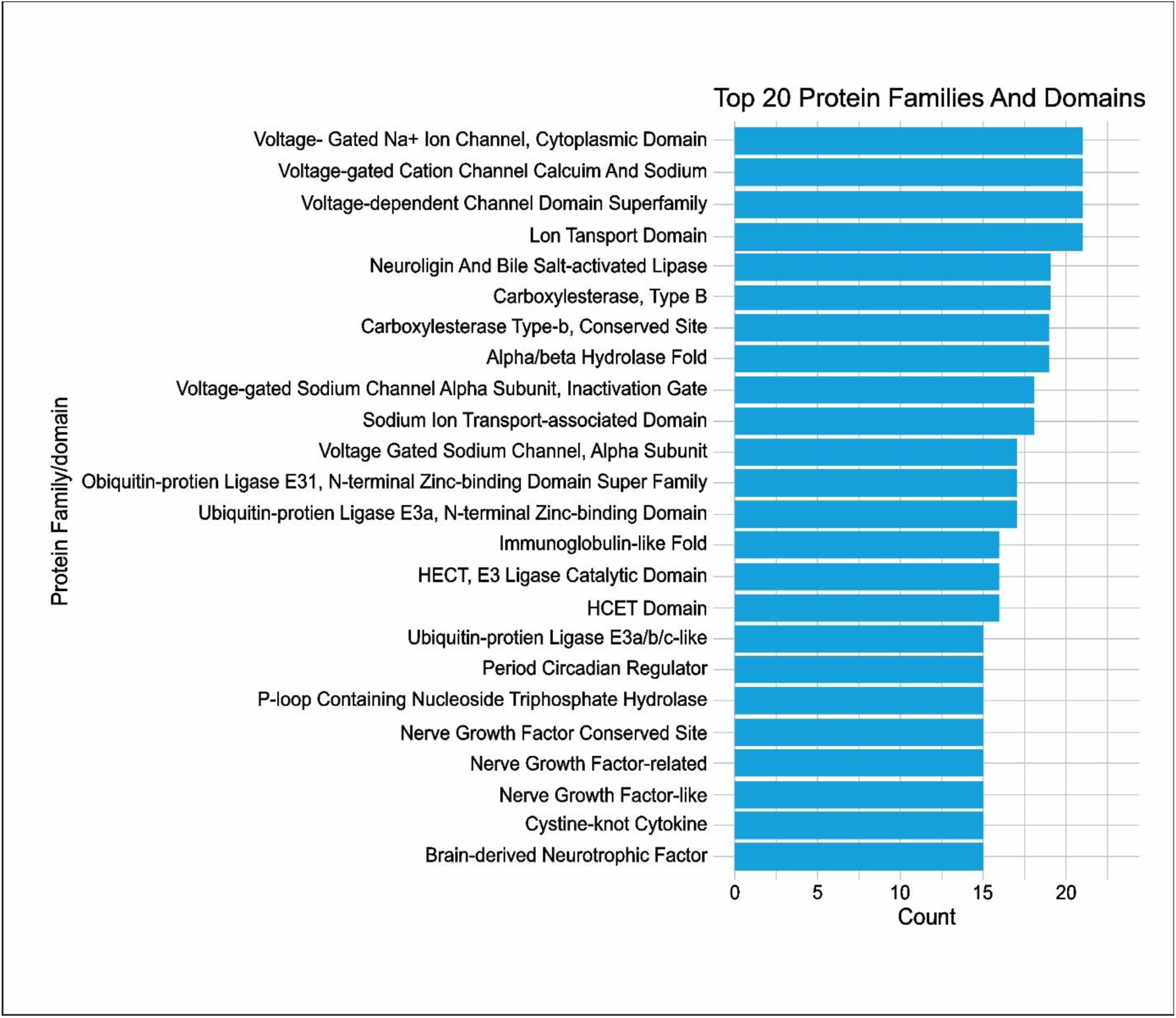
Top 20 enriched protein families and domains. The x-axis represents the count of proteins associated with each family or domain, while the y-axis lists the protein families and domains. Key enrichments include ion channels, metabolic enzymes, ubiquitin-related domains, and neurotrophic factors, reflecting roles in signaling, metabolism, protein regulation, and neuronal function.

Names such as “Lon Transport Domain” and “Neurolysin and Bile salt-activated lipase (BAL)” indicate that the protein may play a role in protein transport and lipid catabolism. The augmentation of COG “Carboxylesterase, Type B” and the PFAM “α/β-hydrolase fold” indicates the existence of analogous enzymes involved in hydrolysis and metabolism, suitable for the breakdown of complex compounds to maintain intracellular homeostasis.

The proteins participating in the ubiquitination process are prominently represented, as indicated by the density of “Ubiquitin-Protein Ligase E3a (UBE3A), N-terminal Zinc-Binding Domain” and associated domains. These domains participate in protein breakdown and signaling via UPS, elucidating regulatory mechanisms governing protein turnover and cellular stress.

Furthermore, the designations such as “Nerve Growth Factor (NGF) Conserved Site,” “Nerve Growth Factor-Related,” and “Brain-Derived Neurotrophic Factor (BDNF)” indicate that neurotrophic signaling pathways pertinent to the survival, growth, and differentiation of neurons are significant to the family. The detected proteins also include the “Period Circadian Regulator” domain, indicating the involvement of circadian clock-associated proteins.

## DISCUSSION

This study aimed to elucidate the evolutionary history of sleep-related gene orthologs, genes with analogous functions across different species. It employed an evolutionary history model focusing on the genes’ origins and evolution rather than their specific functions within individual organisms. This work proved that sleep pathways exhibit evolutionary conservation and found the LUCA genes of human sleep genes. Our phylogenomic approach excludes the physiological, behavioral, or metabolic characteristics related to sleep in LUCA or any other examined animals. Instead, it provides information on the maintenance and advancement of sleep circuits. This study’s rationale was based on the work of Li et al. [24] and aimed to enhance phylogenetic research by utilizing genome-level data, a characteristic of phylogenomics [25–27]. Consequently, whereas traditional phylogenetics often analyzes only 2-3 genes, phylogenomics employs whole-genome data to investigate evolutionary processes. This study is among the rare instances employing this strategy in sleep research.

The findings of the current investigation indicate a significant conservation of sleep-related genes. Several genes, including glutamine synthetase, MHC class II antigens, and Sirtuin 1, are highly conserved, exhibiting an identity exceeding 99%, and thus are deemed physiologically significant. Conversely, the proteins Titin and Hypoxia-inducible factor 1-alpha (HIF-1α) exhibit reduced conservation, suggesting potential adaptive or interspecies variations. The finding of conserved molecular chaperones, including DnaK, casein kinases, and voltage-dependent calcium channels, links them to critical physiological functions such as homeostasis, circadian rhythm, and signal transduction, respectively. BLAST analyses of sleep gene orthologs in Lokiarchaeota, *Chlamydomonas reinhardtii*, *Sulfurimonas paralvinellae*, and LUCA demonstrate that sleep genes and pathways are conserved and suggest potential novel applications of these processes. The highly conserved genes encompass the molecular chaperone DnaK, casein kinase I, and voltage-dependent calcium channels, which have roles in cellular homeostasis, stress response, circadian rhythm, and signal transduction pathways. Uncharacterized proteins in *C. reinhardtii* and bifunctional serine/threonine kinases in *S. paralvinellae* suggest that sleep-like activities are subject to selective pressures for adaptation to environmental or metabolic demands [28,29]. The conserved metabolic enzymes ATP citrate synthase and glutamine synthetase across various taxa underscore the biological importance of sleep in the control of energy homeostasis and cellular repair, pointing to the evolutionary significance of sleep.

The Gene Ontology (GO) enrichment analysis indicated that sleep-related genes are implicated in nervous system development, circadian rhythm regulation, and activity of the cardiovascular and muscular systems. Examining molecular function also emphasized their importance in transcription regulation, ion channel activity, and epigenetic regulation. They suggest that these sleep genes govern nearly all physiological systems in the body, including brain plasticity, cognitive processes, and cardiovascular health. As determined by the DAVID software, the functional classifications of the genes highlighted their importance in pathways, including cAMP signaling, circadian rhythm regulation, and immune system activity [30].

This study revealed the significant evolutionary conservation of sleep/consolidation-related genes; some methodological limitations and alternative causes should be noted. Using bioinformatics techniques, such as BLAST and Gene Ontology enrichment analysis, introduces the possibility of annotation errors, database biases, and computational artifacts that may influence the results. The finding of comparable genes in LUCA and other ancient species does not imply that these genes were initially engaged in sleep control. Their conservation may be attributed to broader cellular activities (e.g., metabolism, stress response, signal transduction) that were subsequently adapted to regulate sleep-related processes in more complex organisms.

This discovery raises intriguing inquiries regarding sleep-related orthologs in LUCA. Initially, it suggests the potential relatedness between humans and flies based on the significance of these genes during the early stages of life. While these genes may have initially served different main functions in organisms, evolutionary influences may have repurposed them for distinct roles in more complex organisms. This phenomenon is termed exaptation, wherein genes are repurposed for a different function at a later period. Nevertheless, caution is necessary when interpreting the findings derived from this study. Sleep-related genes in LUCA do not imply that LUCA experienced sleep or associated events. These genes likely possessed more rudimentary tasks related to cellular functions such as signaling or maintenance, which then evolved into specialized functions in more advanced species. Moreover, contamination or misinterpretation during bioinformatics analyses is unlikely to be eliminated as a source of potential artifacts; thus, validation is essential.

A single gene can influence multiple phenotypes, complicating the association of conserved genes with specific human traits or disorders. Within the context of evolution, pleiotropy suggests that comparative genomics, functional tests, and genome-wide association studies (GWAS) are essential for functional characterizing genes across evolutionary history. Exaptation complicates conclusions, as genes may have originated for one function but were repurposed for sleep-related tasks. Pleiotropy suggests that these genes may influence several physiological pathways beyond sleep, challenging the attribution of their evolutionary conservation solely to sleep-related functions. Future research integrating functional genomics, experimental validation, and comparative phylogenetic methods will be essential for enhancing our comprehension of the evolutionary origins and roles of sleep-related genes.

## CONCLUSION

This is the first-ever study to attempt a clarification of the evolutionary conservation of sleep-related gene orthologs by a phylogenomic methodology. We demonstrated that the genes and processes related to sleep from the reconstructed genome of LUCA are preserved, indicating that sleep is as ancient as life itself. While these genes may have originated and evolved to address fundamental cellular functions, the theory of genetic evolution posits that they may, over time, be repurposed for specialized roles in the control of sleep and circadian rhythms in more complex species. The findings indicate that sleep-related pathways are primordial and contribute to essential ontogenetic processes, including nervous system development, signal transduction, and metabolic functions. Since most of these genes are conserved among several species, they are crucial for cellular and systemic regulation. However, these results should be approached cautiously, as LUCA’s correlation between these genes and sleep does not imply that LUCA slept.

This study examines the issues of evolutionary pleiotropy when genes influence many attributes across several species and evolutionary stages. Future research employing molecular phylogenetics, functional characterization, and comparative analyses is essential to elucidate the regulatory roles of these conserved genes in sleep-specific functions in humans.

Our data extends to LUCA, and although numerous contemporary sleep genes possess conserved homologs in LUCA, this does not imply that sleep is a direct inheritance or as ancient as life itself; thus, we have exercised caution in discussing our findings and conclusions. Our data point to the widespread role of genetic conservation in biological activity; nevertheless, we do not provide concrete proof that LUCA or any primordial living forms exhibited sleep-like behaviors. Instead, these conserved genes may have initially fulfilled vital cellular functions that were subsequently repurposed for sleep control in more complex organisms. Future experimental research is required to validate these genes’ functional involvement in sleep processes and the broader physiological importance of the proposed pathways throughout evolution.

## CRediT authorship contribution statement

SRP; SBC: Conceptualization and Methodology; SRP; KMS; SP: Data collection and review, KMS; SP: Software, SRP; KMS; SP: Preparation of original draft; DWS; SBC; GPN: Critical review and editing; SBC: supervision; Prior to submission, all authors (SRP; KMS; DWS; SP; GPN, SBC) have read and agreed to the submitted version of the manuscript.

## Author Agreement Statement

All authors consciously confirm that the paper is their original work that has not previously been published or is not presently being considered for publication anywhere. Each author expressly declares that they contributed equally, read, evaluated, and accepted the final version of the paper, and agreed to be included as co-authors per ICMJE guidelines. Each of the authors has approved the author sequence and the corresponding authors.

## Role of medical writer or editor

Not applicable.

## Ethical Statement

This study does not contain any work involving animals or human participants performed by any authors. Hence, no IRB approval was necessary for this work.

## Data availability

This study utilized the dataset that was originally prepared for our ongoing phylogenomics investigation of sleep. Since this is a work in progress, no additional information will be available from the authors.

